# Male mice heterozygous for Protamine-1 and Protamine-2 are infertile displaying sperm damage and retention of Protamine-2 precursors, transition proteins and histones

**DOI:** 10.64898/2026.03.15.711850

**Authors:** Gina Esther Merges, Christoph Wiesejahn, Mar Domingo-Lopez, Simon Schneider, Andjela Kovacevic, Lena Arévalo, Hubert Schorle

## Abstract

**BACKGROUND:** During spermiogenesis, histones are exchanged by protamines (*PRMs*) in spermatids, which results in DNA hypercondensation and protection. Rodents and primates express two *PRMs* (*PRM1* and *PRM2*) in a species-specific ratio. Maintaining this ratio is necessary for functional chromatin reorganization and alteration is associated with sub- or infertility in mice and humans. *Prm1* and *Prm2* deficient mice are infertile, while *Prm1*^*+/-*^ males are subfertile showing a severely altered PRM ratio. *Prm2*^*+/-*^ males are fertile and display a protamine ratio comparable to WT.

**OBJECTIVES:** Here, we addressed the question whether loss of one allele of *Prm1* and one allele of *Prm2* affects fertility.

**MATERIAL AND METHODS:** Double heterozygous (dHET) mice lacking one allele of *Prm1* and one allele of *Prm2* were generated and analyzed

**RESULTS:** dHET males were infertile with sperm showing retention of histones and TNPs, high levels of PRM2 precursor and decreased levels of mature PRM2. In mature sperm the PRM ratio and the total PRM content was not altered. However, CMA3 staining revealed incomplete protamination and sperm nuclei appeared more rounded and slightly bigger, suggesting impaired DNA-hypercondensation. In dHET sperm, DNA degradation was apparent, but to a lower level compared to sperm from *Prm1* and *Prm2* deficient males. Increased 8-OHdG levels suggested oxidative stress in the epididymis of dHET mice. However, a fraction of dHET sperm were capable of fertilization, with embryonic development up to 8-cell stage.

**DISCUSSION AND CONCLUSION:** These results suggest, that male factor infertility might not be reliably detected by measuring PRM1/PRM2 ratio but rather by determining the level of protamination by e.g. CMA3 analysis and pre-PRM2 retention.

## Introduction

Sperm require a strong reduction in nuclear size and increased measures of chromatin protection in order to safely and effectively migrate through the female reproductive tract. During late spermiogenesis, the majority of somatic histones are replaced by sperm-specific histone variants and transition proteins (TNP1 and TNP2) ^1-4^. Protamines then replace histone-TNP complexes ^2, 5-7^. Upon completion of protamination, the DNA is hypercondensed, the paternal genome protected and transcription silenced ^8, 9^. Protamine 1 (*Prm1/PRM1*) is present in sperm of all mammalian species, while primates and rodents additionally express protamine 2 (*Prm2/PRM2*). Both genes (*Prm1* and *Prm2*) map within 12 kb on chromosome 16 in humans and mice ^10, 11^. *PRM2* is expressed as a precursor protein. Upon binding, the N-terminal domain (cP2) is cleaved off (processed) giving rise to mature PRM2 (mP2) which mediates DNA hypercondensation in conjunction with PRM1 ^5, 12^. PRM1 and PRM2 are found in a species-specific ratio (1:1 in humans ^13^; 1:2 in mice ^14^). Alterations of the ratio and reduction of the protamine amount are correlated to sub- and infertility ^15-26^.

We have shown that loss of *Prm1* or *Prm2* (*Prm1*^*-/-*^ or *Prm2*^*-/-*^) led to male mice infertility ^26-28^. Sperm were inviable, immotile and accumulated reactive oxygen species (ROS)-mediated damage. Moreover, histone and transition protein (TNP) retention was observed in both *Prm1*^*-/-*^ and *Prm2*^*-/-*^ sperm.

Here we generated *Prm1*^+/-^*Prm2*^*+/-*^ double heterozygous (dHET) mice and found the loss of two Protamine alleles did not affect overall Protamine content in dHET sperm. However, male dHET mice were infertile, with sperm displaying retained histones, TNPs and pre-PRM2. CMA3 staining revealed incomplete Protamination to an even higher extent than seen in the single *Prm* knockout mouse models. While sperm viability and motility were significantly reduced and epididymal sperm DNA was fragmented, in contrast to *Prm1*^*-/-*^ and *Prm2*^*-/-*^ males, some dHET sperm appeared to be capable of fertilization with developmental arrest of embryos up to the 8-cell stage.

## Materials and Methods

### Ethics statement

Animal experiments were conducted in accordance to the German law of animal protection and approved by the local institutional animal care committees (Landesamt für Natur, Umwelt und Verbraucherschutz, North Rhine-Westphalia, approval ID: AZ84-02.04.2013.A429; AZ81-0204.2018.A369).

### Generation of *Prm1 Prm2* double heterozygous mice

*Prm2*^*-/-*^ females of the *B6-Prm2197* line ^27^ were mated with C57BL/6J WT males to obtain *Prm2*^*+/-*^ zygotes. Zygotes were isolated 0.5 dpc and electroporated with guide RNAs targeting *Prm1* in exon 1 and exon 2 and the Cas9 endonuclease. The guides were in sequence equivalent to those used to generate *Prm1*-deficient mice ^26^. Next, zygotes were transferred to pseudo-pregnant foster mice. Founder animals were sequenced and a mouse carrying a deletion equivalent to the B6-*Prm1Δ167* line was selected for mouse line establishment. Additionally, a line was established from a mouse carrying a 139 bp deletion in the *Prm1* coding sequence. Mice were genotyped using the *Prm1*- or *Prm2*-PCRs described before ^26, 27^.

### Mouse genotyping and sequencing

Mice were genotyped as described before ^26, 27^. Primers flanking the gene-edited region were used (*Prm2*_fwd: 5’-TGCAGCCTCAATCCAGAACC, *Prm2*_rev: 5’-TGTAGCCTCTTACGAGAGCAG; *Prm1*_fwd: 5’-CCACAGCCCACAAAATTCCAC, *Prm1*_rev: 5’-TCGGACGGTGGCATTTTTCA). Cycling conditions for *Prm2* were: 2 min 95 °C; 30x (30 s 95 °C; 30 s 60 °C; 30 s 72 °C) 5 min 72 °C. Cycling conditions for *Prm1* were: 2 min 95 °C; 35x (30 s 95 °C; 30 s 64 °C; 30 s 72 °C) 5 min 72 °C. For *Prm2* amplification of the WT allele resulted in a 220 bp product and amplification of the *Prm2Δ97* allele in a 123 bp product. Amplification of the *Prm1* locus produced a 437 bp product from the WT allele and products of 298 bp and 270 bp from the *Prm1Δ139* and *Prm1Δ167* allele, respectively.

For validation by sequencing, PCR products were cloned into the pCR 2.1-TOPO plasmid (Thermo Fisher Scientific) following the manufacturers recommendations. Plasmids were transformed into E.cloni 10G Chemically Competent Cells (Lucigen), isolated by alkaline lysis and sequenced (GATC/Eurofins).

### Fertility test

Male mice at an age of 8 to 13 weeks were bred 1:1 or 1:2 with C57BL/6J females to assess fertility. Females were aged between 10 and 16 weeks and checked daily for the presence of seminal plugs. Plug-positive females were separated from the males and examined for pregnancies and litter sizes. At least 5 plug-positive females were monitored per male.

### Tissue and sperm sampling

Animals were sacrificed by cervical dislocation and weighed. Epididymides were dissected, and testes were dissected and weighed. For mature sperm isolation, the cauda epididymis was dissected and cut several times in M2 medium or PBS at 37°C. After 15 min incubation time, tissue pieces were gently rinsed, assuring residual sperm leave the cauda. Testicular spermatids were isolated as described by Kotaja *et al*. ^29^. Using the Leica MS5 stereomicroscope and Schott KL1500 light source, testes were decapsulated and gently pulled apart while immersed in PBS to release single seminiferous tubules. Elongating and condensed spermatid containing sections were identified through their light absorption pattern. Selected tubule sections were placed on a microscope slide. By adding a coverslip, the tissue was squashed, pushing out spermatogenic cells. Samples were frozen in liquid nitrogen, the coverslip was flipped off, the slide was immersed in 90% EtOH for 5 min and dried. Tissues and isolated sperm were processed depending on the required experiment (see below).

### Macroscopic analysis of testis

Sections of Bouin-fixed testis were processed and stained with hemalum solution acid ^30^ and Eosin Y solution (Carl Roth) as described previously ^26^.

### Immunohistochemistry

Bouin’s solution (75% saturated picric acid, 10 % formalin, 5 % glacial acetic acid) or paraformaldehyde (PFA; 4 % w/v PFA/water) were used to fix tissue samples (4°C, overnight). Tissues were embedded in paraffin and cut into 3 µm sections using the Microm CP60 microtome. Samples were deparaffinized, rehydrated and treated with DNA decondensation buffer [0.385 g DTT, 200 μl Triton X-100, 200 i.u./ml heparin, ad 100 ml PBS]. Antigen retrieval was done by boiling for 15 min in citrate buffer (pH 6), followed by blocking in horse serum. Primary antibodies were diluted in 5% BSA in PBS and applied to the sections, followed by incubation overnight at 4°C [8-OHdG (abcam; ab48508; 1:500), cP2 (Arévalo *et al*. 2022, Davids Biotechnologie GmbH; 1:2500), histone H3 (abcam; ab1791; 1:1500), TNP1 (abcam; ab73135; 1:1000), TNP2 (Santa Cruz; sc-393843; 1:100)]. After washing in PBS, sections were incubated with the secondary antibodies for 1 h at room temperature [Vector Laboratories; Vectafluor Kits; DI-1788, DI-1794, DK-8828]. Finally, samples were washed again and counterstained using ROTI Mount FluorCare DAPI. Imaging was performed using a confocal Visitron VisiScope and the VisiView Software or a Leica DM5500 B microscope.

### MitoTracker Red – PNA-FITC staining

Sperm fixed in 4% PFA were washed in PBS and incubated in MitoTracker Red CMXRos (25 nM, Cell Signaling Technology) and Lectin PNA Conjugates (0.02 mg/ml, Molecular Probes) for 45 min. Cells were washed again and dropped onto slides. After drying, sperm were counterstained using ROTI Mount FluorCare DAPI.

### Transmission Electron Microscopy

TEM processing and imaging was performed by the University of Bonn Imaging core facility, as described before ^31^. Pelleted sperm and testicular tissue were fixed in 3% glutaraldehyde overnight at 4 °C. Then they were washed, post-fixed with 2% osmium tetroxide for 2 h at 4°C and washed again. The samples were dehydrated in an ascending ethanol series and contrasted in 70% (v/v) ethanol with 0.5% uranyl acetate for 1.5 h at 4°C. Afterwards, they were washed with propylene oxide and stored in propylene oxide:Epon C (1:1) at 4°C overnight. The pellets were embedded in Epon C and ultrathin sections were prepared. The samples were contrasted again with uranyl contrasting solution and lead citrate. Imaging was carried out with a Zeiss Crossbeam 550 equipped with a STEM detector using 20-25 kV and 100-150 pA. Zeiss Atlas 5 and SmartSEM software were used for image processing.

### Assessment of sperm DNA fragmentation

Genomic DNA was isolated from frozen sperm according to a previously described protocol with small adjustments ^32^. Sperm were incubated in lysis buffer [1 M Tris-HCl pH 8.0, 3 M NaCl, 0.5 M EDTA, 20% (m/v) SDS] at 50°C overnight with supplementation of 21 μl 1 M DTT, 2.5 μl 0.5 % Triton X-100 and 40 μl 10 mg/ml proteinase K. Samples were centrifuged (15.500 g, 10 min) and the supernatant was mixed with 1 μl 20 mg/ml glycogen and 1/10 vol 3 M NaAc. DNA was precipitated with 100% ethanol for at least 2 h at -80°C and the pellet was washed with 75% ethanol. It was dried using a Speed Vac DNA110 (Savant) and dissolved in TE buffer. DNA was separated on a 0.8% agarose gel. Part of the results have been published before ^26^.

### Chromomycin A3 staining

Mature sperm were fixed in Carnoy’s solution [3:1 methanol:acetic acid (v/v)] after swim-out, spread out and dried on microscope slides. Samples were covered with 100 μl CMA3 solution [0.25 mg/ml CMA3 in Mcllvaine buffer (pH 7.0, containing 10 mM MgCl2)] and incubated for 20 min in the dark. Then, slides were washed with PBS and counterstained and mounted with ROTI Mount FluorCare DAPI.

### Viability test

50 μl of sperm swim-out was thoroughly mixed with 50 μl of EN staining solution (Eosin-Nigrosin, Morphisto, Offenbach, Germany). 30 μl of the suspension was smeared on microscope slides followed by mounting with Entellan.

### Morphological analysis of sperm nuclei

Epididymal sperm fixed with Carnoy’s solution were dropped onto a slide, dried and mounted with DAPI containing medium (ROTI Mount FluorCare DAPI). Analysis was carried out using the software ‘Nuclear_Morphology_Analysis_1.18.1_standalone’ ^33^. A minimum of 100 sperm per mouse and three mice per genotype were analyzed. The shape of step 15-16 testicular sperm was analyzed from DAPI stained smashed tubule preparations. A minimum of 250 spermatids per sample were analyzed. Pairwise comparisons were calculated by the Mann-Whitney U test and p-values were corrected using Bonferroni correction.

### Sperm motility analysis

For motility assessment, sperm swim-out from cauda epididymidis was done using Gynemed SpermAir medium and samples were incubated for 30 min at 37 °C. Sperm were further diluted 1:20 in prewarmed medium and 30 μl were pipetted onto a slide. A coverslip with a spacer was added and the slide was placed on a heated slide holder at 37 °C. Using the a brightfield microscope (Leica DM-IRB) equipped with a Basler acA1920-155ucMED camera, five 3-second videos were recorded for each sample. Motile sperm were distinguished and counted in ImageJ. At least 200 sperm were monitored per individual for 3 individuals per genotype.

### Sperm basic protein extraction

The isolation of protamines and other basic proteins was conducted according to ^34^. Frozen sperm samples were rinsed in PBS and resuspended in buffer [4 μl 1 M Tris pH 8, 0.8 μl 0.5 M MgCl2, 5 μl Triton X-100]. Testis tissue was homogenized and filtered through a 35 µm cell strainer before adding the buffer. The suspension was centrifuged at 9000 g for 5 min at 4°C and 1 mM phenylmethylsulfonyl fluoride (PMSF) was added. Further, solution 1 [10 mM PMSF, 2 mM EDTA, 100 mM Tris, pH 8] followed by solution 2 [0.04435 g DTT in 0.5 ml 6 M GuHCl] were mixed with the lysed sperm samples. Vinylpyridine was added to a final concentration of 0.8% and the samples were incubated for 30 min at 37°C and vortexed every 5 min. 5× ice-cold ethanol (100%) was added to precipitate DNA, proteins dissolved in 0.5 M HCl and precipitated with 100% trichloroacetic acid (TCA) at 4 °C. Proteins were washed twice with acetone, dried completely and resuspended in sample buffer [5.5 M urea, 20% 2-mercaptoethanol, 5% acetic acid].

### Acetic acid-urea / thiourea polyacrylamide gel electrophoresis

Basic protein extractions were separated by acid-urea gel electrophoresis (AU-PAGE) as described by Soler-Ventura *et al*. ^34^. A pre-electrophorized 15 % AU-gel [2.5 M urea, 0.9 M acetic acid, 15% acrylamide / 0.1% N,N′-methylene bis-acrylamide, tetramethylethylenediamine, ammonium persulfate] was used. Alternatively, thiourea gels [2.5 M urea, 12.5 mM thiourea, 0.9 M acetic acid, 15% acrylamide / 0.1% N,N′-methylene bis-acrylamide, 0.2% H2O2] were used without pre-electrophoresis. Bands were either visualized by Coomassie Brilliant Blue staining or used for western blotting.

### Western blots

After separation by AU- or thiourea PAGE, proteins were transferred onto a PVDF membrane using the Bio-Rad Trans-Blot Turbo System. The membrane was blocked using 5% milk powder in TBST (TBS, 0.1% Tween 20) for 1 h at room temperature or overnight at 4 °C. Primary antibodies diluted in blocking solution were added to the membrane and incubated overnight at 4 °C [PRM1 (Briar Patch Biosciences; Mab-Hup1N-150; 1:1000), PRM2 (Briar Patch Biosciences; Mab-Hup2B-150; 1:1000), ODF2 (proteintech; 12058-1-AP; 1:500-1:1000), histone H3 (abcam; ab1791; 1:1000), histone H4 (abcam; ab177840 ; 1:500)]. Then, the membrane was washed in TBST and incubated for 1 h at room temperature with the secondary antibody [Agilent Technologies Singapore; polyclonal goat anti-rabbit IgG/HRP (P044801-2; 1:2000), polyclonal rabbit anti-mouse IgG/HRP (P026002-2; 1:1000)]. The membrane was washed again and protein bands were detected using WESTAR NOVA 2.0 chemiluminescent substrate (Cyanagen) using the Bio-Rad ChemiDoc MP Imaging system. Protein bands were quantified by ImageJ.

### Coomassie stained and WB band quantification

Coomassie Brilliant blue stained thiourea gels were imaged and band sizes were quantified using ImageJ 1.54g software as described previously ^26, 35^. Relative protamine amounts were normalized against total protein levels and total PRM levels.

### Monitoring embryonic development

dHET and WT males were bred with superovulated WT females. After being plug-positive, females were sacrificed on day 0.5 and fallopian tubes were removed and flushed with embryo culture medium (EMD Millipore EmbryoMax M2 medium) to extract oocytes. The cumulus cell layer surrounding oocytes was digested with 0.5 mg/ml hyaluronidase in M2 medium. Then, oocytes were washed several times in M2 medium and deposited in a microdrop of medium (Vitrolife G-TL) in order to monitor further development. The drop was overlayed with Vitrolife embryo grade mineral oil to avoid evaporation of medium.

### Statistical analysis

Values are depicted as means ± standard deviation (s.d.) unless mentioned differently. Student’s t-test or ANOVA were used to calculate statistical significances and a value of P<0.05 was considered significant (*P<0.05; **P<0.005; ***P<0.001).

## Results

### Establishment of mouse line heterozygous for *Prm1* and *Prm2*

PRM1 and PRM2 are highly basic proteins, containing arginine-rich DNA binding domains. To generate mice deficient for both *Prm1* and *Prm2*, previously established *Prm2*^*-/-*^ female mice carrying a 97 bp deletion in the *Prm2* coding sequence ^27^ were superovulated and bred with WT males. The resulting *Prm2*^*+/-*^ zygotes were electroporated with guide RNAs targeting exon 1 and exon 2 of the *Prm1* coding region together with the Cas9 endonuclease. The guide RNAs were identical to those used to generate *Prm1*-deficient mice ^26^. Two mice carrying deletions in both genes on the same allele (in *cis*) were identified. One carried a 167 bp deletion in the *Prm1* coding sequence, leading to loss of the central DNA-binding domains of PRM1 (*Prm11167Prm2197*) (Fig. 1A-B). The *Δ167* deletion in *Prm1* was equivalent to a previously described and analyzed *Prm1*-null allele ^26^. The other carried a 139 bp deletion in the *Prm1* coding sequence, leading to a frameshift and premature stop codon (*Prm11139Prm2197*). Offspring was genotyped (Fig. 1D). For *Prm1* amplification of the wild-type (WT; *+/+*) allele generated a product of 437 bp and the gene-edited alleles *Prm1Δ167* and *Prm1Δ139* of 270 bp and 242 bp, respectively. For *Prm2*, the WT allele resulted in a product of 220 bp, while the edited locus had an amplicon of 123 bp. In addition, double heterozygous male mice carrying the alleles in *trans* were generated by mating *Prm1*^*+/-*^ and *Prm2*^*+/-*^ mice.

**Fig. 1.**
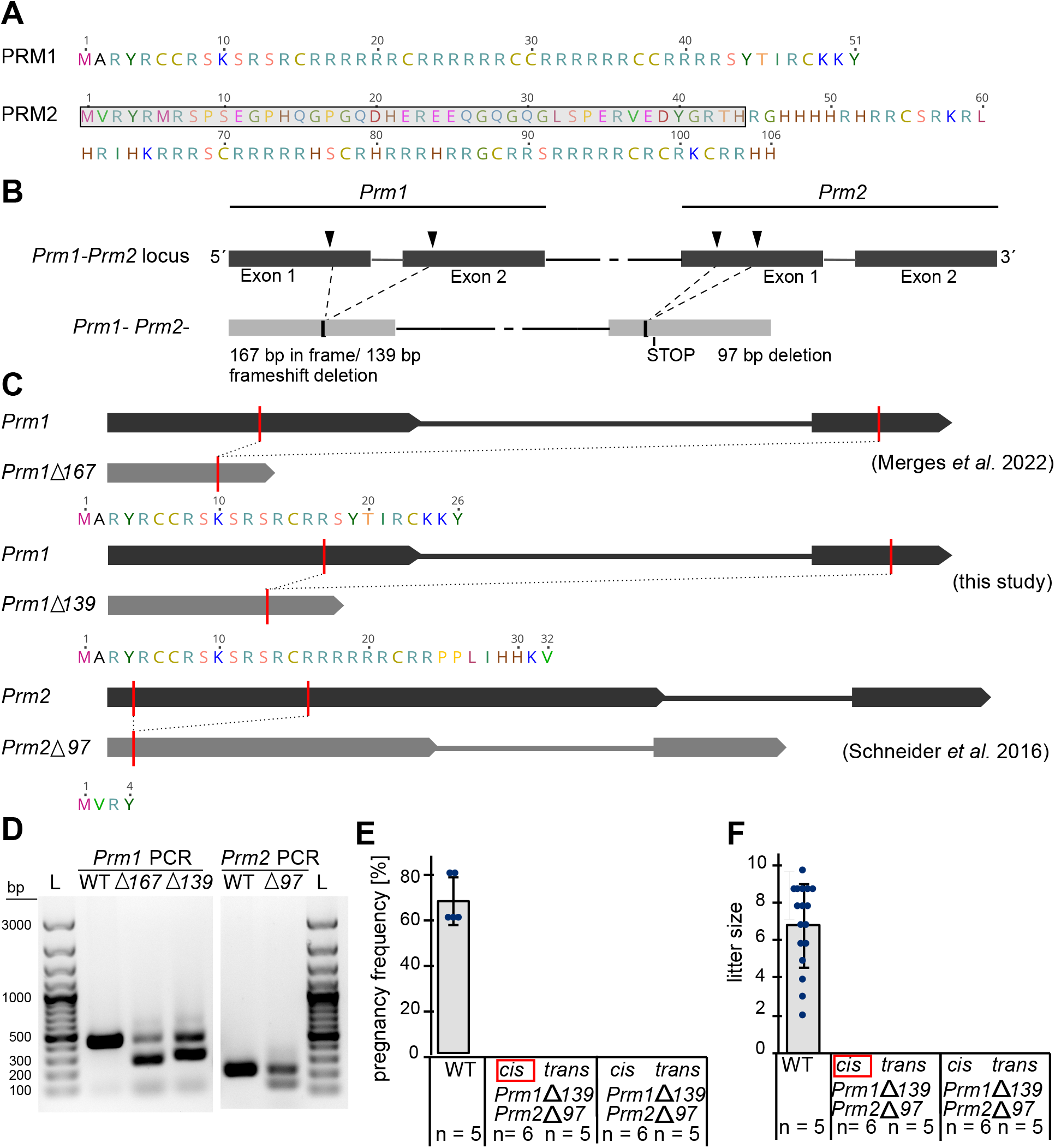
Generation of *Prm1* and *Prm2* gene-edited alleles and male mice fertility. (A) Schematic depiction of murine PRM1 and PRM2 amino acid sequences (grey box = cP2). (B) Schematic depiction of CRISPR/Cas9-mediated gene editing of the *Prm1-Prm2* locus. Black arrow heads indicate the target sites of CRISPR-Cas9 guide RNAs. (C) Schematic depiction of *Prm1* and *Prm2* gene-edited alleles and their respective amino acid predictions. (D) Genomic WT and dHET DNA amplified with PCRs targeting either *Prm1* or *Prm2*. L = ladder. (E) Average pregnancy frequency after mating WT and dHET males with WT C57BL/6J females. Dots represent different males (n = number of males). (F) Average litter size after mating WT and dHET males with WT females. Dots show each litter size (n = number of males).

### Males heterozygous for *Prm1* and *Prm2* are infertile

*Prm2*^+/-^ males were shown to be fertile, while *Prm1*^*+/-*^ males are subfertile ^26, 27^ (Table S1). To test fertility, five (cis) or six (trans) *Prm1*^*+/-*^*Prm2*^*+/-*^ males were used (Fig. 1E-F). No pregnancies were reported by mating males of any of the four *Prm1*^*+/-*^*Prm2*^*+/-*^ genotypes. WT males generated offspring with a pregnancy frequency of 68% (Fig. 1E) and an average litter size of seven (Fig. 1F). For further analyses males harboring the *Prm11167Prm2197* mutations on one allele (in cis) were used and are henceforth referred to as dHET.

### dHET males show no changes in protamine ratio or total protamine content but large amounts of retained PRM2 precursors

Since dHET males lack one copy of each protamine, we expected the total protamine content to be reduced but protamine ratio to be unaltered. Both PRM1 and PRM2 were detected using immunohistochemical staining on testis sections (Fig. S1). To analyze the state of protamination in dHET cauda epididymal sperm, we first examined the PRM ratio and content by acid urea PAGE (AU-PAGE) and thiourea PAGE (Fig. S2). Since protamine bands separate better on thiourea gels, these were chosen for analysis (Fig. 2A). Surprisingly, the total PRM content calculated by normalizing the PRM signal against the whole protein per lane showed no significant difference between WT (mean: 21.24%) and dHET (mean: 19.96%) (Fig. 2B). Also, the total PRM2 and PRM1 levels relative to total protein did not differ significantly between dHET and WT samples (Fig. 2C-D). However, while all detectable PRM2 was fully processed to mP2 in WT sperm, we found large amounts of PRM2 precursors in dHET samples (Fig. 2A dashed box). Relative levels of mP2 were significantly reduced in dHET sperm compared to WT sperm (Fig. 2E), while PRM2 precursor forms were retained (Fig. 2F). Noteworthy, an additional band was prominent in dHET samples (Fig. 2A red arrow), suggesting a partially processed precursor. Lower levels of mP2 and higher levels of PRM2 precursor were also significantly different when being compared to the total PRM levels (Fig. 2G-H). The PRM1:PRM2 ratio did not significantly differ considering total PRM2 including PRM2 precursors (Fig. 2I). Pre-PRM2 accounted for 60.36% of total PRM2 in protein extraction from dHET sperm (Fig. 2J).

**Fig. 2.**
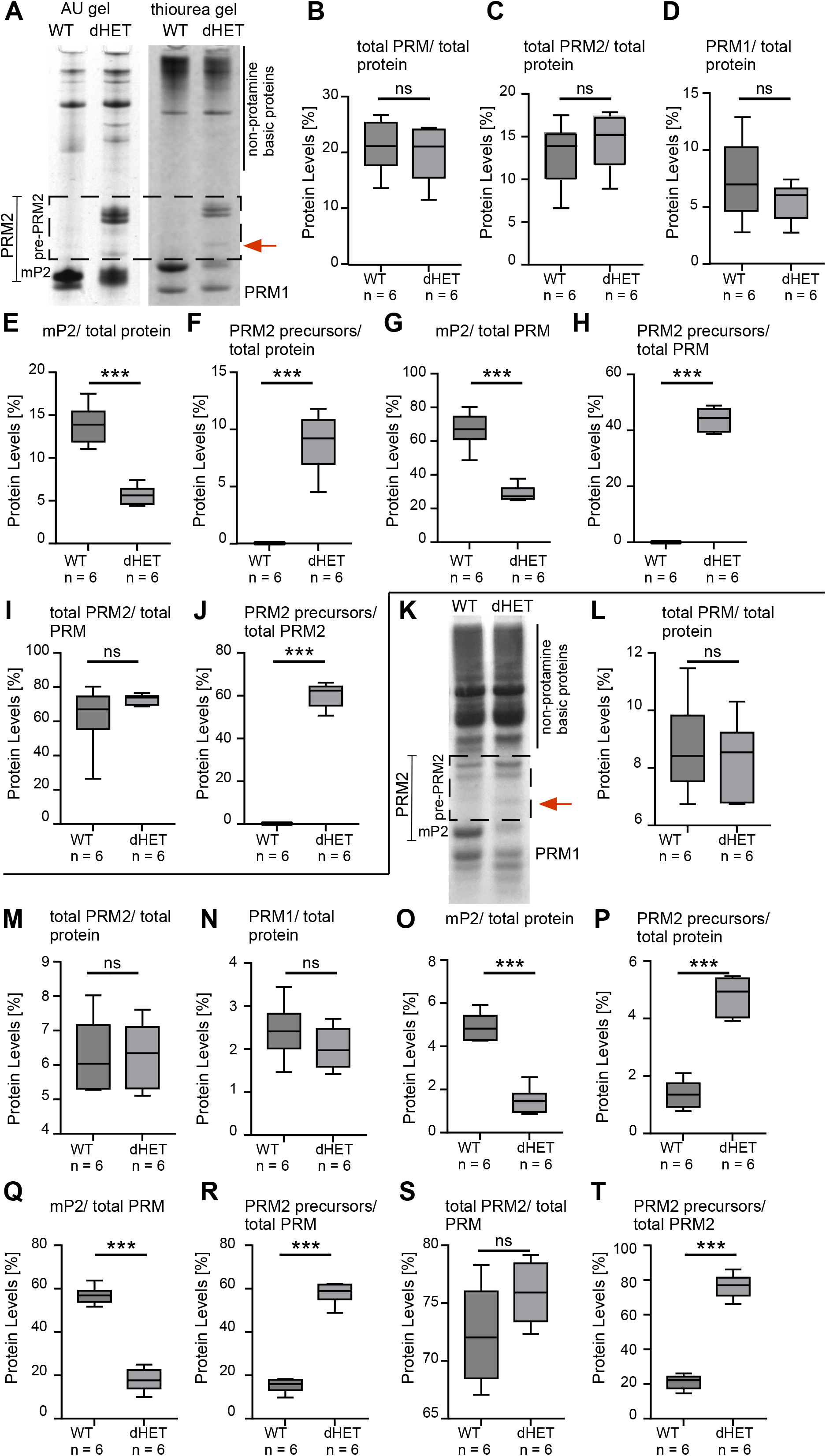
Protamine content and ratio in dHET mice. (A) Separation of basic cauda epididymal sperm protein extractions on Coomassie-stained acid-urea (AU) and thiourea gels. PRMs are detected at the bottom of the gel. Precursor bands of PRM2 are prominent in dHET samples (dashed box). Thiourea gels were used for analyses shown in (B-I). (B) Relative level of protamines compared to total protein content extracted from WT and dHET sperm. (C) Relative level of total PRM2 compared to total protein content extracted from WT and dHET sperm. (D) Relative level of PRM1 compared to total protein content extracted from WT and dHET sperm. (E) Relative level of mP2 compared to total protein content extracted from WT and dHET sperm. (F) Relative level of PRM2 precursors compared to total protein content extracted from WT and dHET sperm. (G) Relative level of mP2 compared to total PRM content extracted from WT and dHET sperm. (H) Relative level of PRM2 precursors compared to total PRM content extracted from WT and dHET sperm. (I) Relative level of total PRM2 precursors compared to total PRM content extracted from WT and dHET sperm. (J) Relative amount of PRM2 precursors comared to total PRM2 extracted from WT and dHET sperm. (K) Separation of basic protein extractions from whole testis on thiourea gels stained with Coomassie. PRM2 precursors are marked with a dashed box. Thiourea gels were used for quantifications shown in (L-S). (L) Relative level of protamines compared to total protein content extracted from WT and dHET testis. (M) Relative level of total PRM2 compared to total protein content extracted from WT and dHET testis. (N) Relative level of PRM1 compared to total protein content extracted from WT and dHET testis. (O) Relative level of mP2 compared to total protein content extracted from WT and dHET testis. (P) Relative level of PRM2 precursors compared to total protein content extracted from WT and dHET testis. (Q) Relative level of mP2 compared to total PRM content extracted from WT and dHET testis. (R) Relative level of PRM2 precursors compared to total PRM content extracted from WT and dHET testis. (S) Relative level of total PRM2 precursors compared to total PRM content extracted from WT and dHET testis. (T) Relative amount of PRM2 precursors comared to total PRM2 extracted from WT and dHET testis.

To explore the protamine content during spermiogenesis, basic proteins were isolated from whole testis and separated on thiourea gels (Fig. 2K, Fig. S3). No significant differences in PRM content compared to total protein were detected in dHET compared to WT samples (Fig. 2L). Similarly, the total PRM2 and PRM1 content was not significantly different from WT samples (Fig. 2M-N). However, significantly lower amounts of mP2 were detected in dHET samples (Fig. 2O) and levels of PRM2 precursors was significantly higher in dHET samples (Fig.2P). Again, elevated levels of PRM2 intermediates were detected in dHET samples (Fig. 2K red arrow). Decreased levels of mP2 and increased levels of PRM2 precursors in dHET compared to WT samples were also significant when compared to the total PRM amount (Fig. 2Q-R). When comparing the total PRM2 level to the total amount of PRM, WT samples showed a higher variability, but differences between dHET and WT were not significant (72.28 ± 4.288 % in WT and 75.89 ± 2.580 % in dHET) (Fig. 2S). The levels of pre-PRM2 relative to total PRM2 were significantly increased in dHET testis compared to WT testis (21.23 ± 4.199 % in WT and 76.47 ± 6.804 % in dHET) (Fig. 2T).

### dHET sperm presented with reduced motility, viability and morphological defects

The testis to body weight ratio was not significantly different between dHET and WT males (Fig. 3A). Spermatogenesis appeared normal in dHET males (Fig. 3B). Sperm viability was reduced in dHET (29% viable) compared to WT (64% viable) sperm (Fig. 3C). In comparison, *Prm1*^*+/-*^ and *Prm2*^*+/-*^ strongly resembled the WT, while *Prm1*^*-/-*^ and *Prm2*^*-/-*^ displayed only 1% and 5% viable cells respectively ^25, 28^. Additionally, only 9% of sperm from dHET mice were motile compared to 71% of sperm from WT mice (Fig. 3D). A large portion of mature dHET sperm showed disruptions of the 9+2 axonemal structures in flagellae (Fig. 3E, S4).

**Fig. 3.**
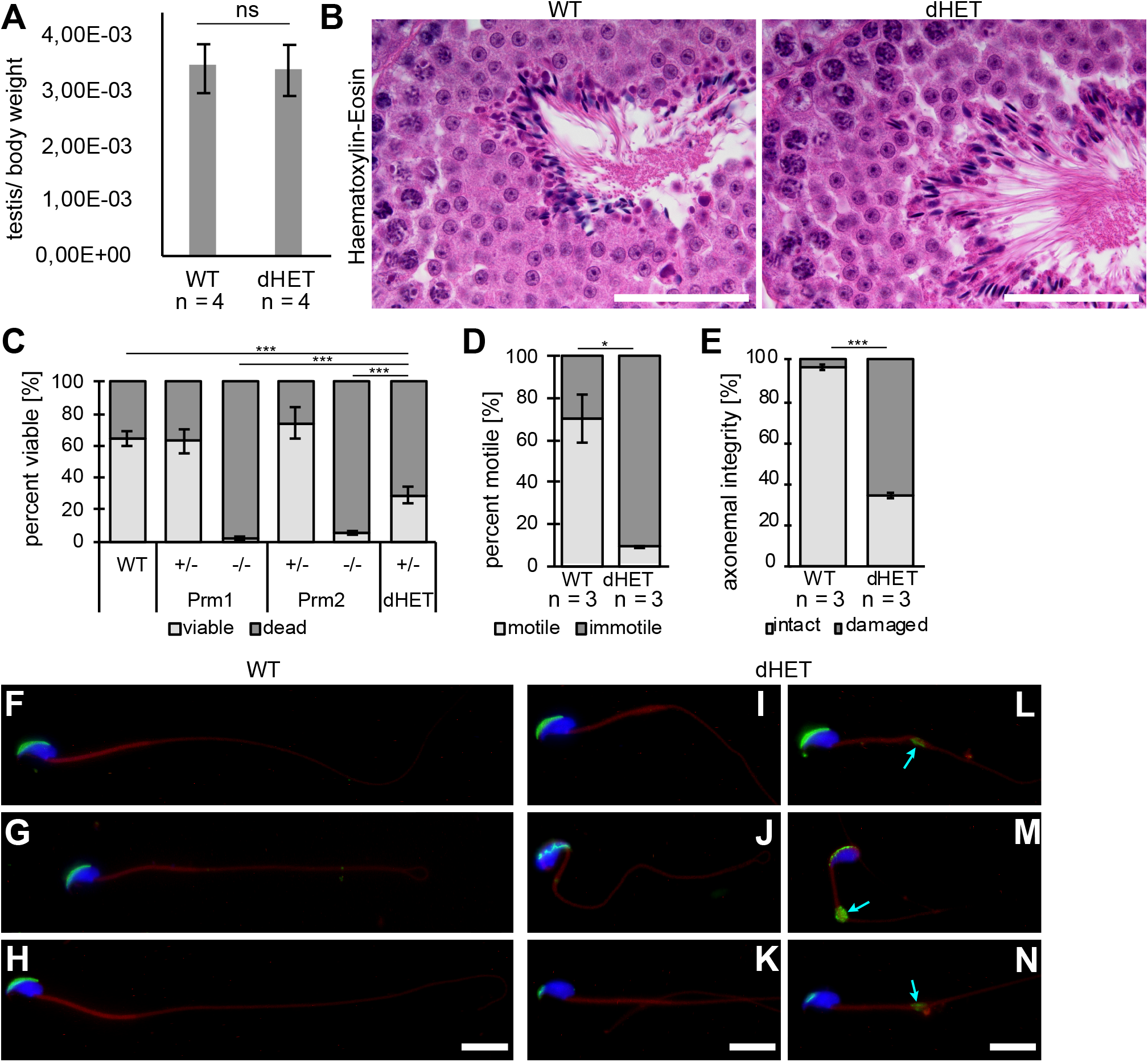
Spermatogenesis and sperm morphology of dHET mice. (A) Testis to body weight ratio of dHET and WT mice (n= number of males). (B) Representative stage VII-VIII seminiferous tubule section of dHET and WT stained with Hematoxylin and Eosin. (C) Percentages of viable and inviable mature sperm of WT, *Prm1*^*+/-*^, *Prm1*^*-/-*^, *Prm2*^*+/-*^, *Prm2*^*-/-*^ and dHET mice. Part of the data has been published before ^26, 28^. Data are mean ± s.d. and was analyzed by ANOVA. (D) Percentages of motile and immotile sperm in WT and dHET mature swim-out samples (n= number of males). (E) Quantification of axonemal integrity. A minimum of 100 flagella cross sections were per animal were analyzed (n= number of males). Data are mean ± s.d. and were analyzed by two-tailed, unpaired Student’s t-test (***p<0.001). (F-N) Sperm stained with DAPI (blue), PNA-FITC (green) and MitoTracker (red). Turquoise arrows highlight excess residual cytoplasm. (C). Scale bars: 50 μm (B); 10 μm (F-N).

Next, we stained mature sperm with PNA-FITC and MitoRed (Fig. 3F-H). In dHET sperm some acrosomes appeared to be intact (Fig. 3I), while others appeared irregular (Fig. 3J, L) or displayed only small residues of acrosomal structures (Fig. 3K, M-N). Of note, we found many sperm in dHET samples that showed excess residual cytoplasm at the distal end of the midpiece, which is known to be an aberrant structure associated with defective spermatogenesis ^36^ (Fig. 3L-M).

Taken together dHET sperm show acrosome, flagella and membrane damage which explains the reduced sperm motility and viability. These defects were more severe compared to *Prm1*^*+/-*^ and *Prm2*^*+/-*^ samples, but not as pronounced as seen in *Prm1*^*-/-*^ and *Prm2*^*-/-*^ mice.

### dHET sperm showed increase in nuclear size, partially fragmented DNA and oxidative stress induced sperm damage

Nuclear sperm head morphology was assessed via DAPI staining. Mature dHET sperm presented as two morphologically different fractions, which we refer to as DAPI-bright and DAPI-weak (Fig. 4A). DAPI-bright cells strongly resemble those from WT, while DAPI-weak, duller sperm are smaller, suggestive of DNA damage. Using an ImageJ plugin, we carried out nuclear morphology analysis of WT and DAPI-bright dHET cauda epididymal sperm. The analysis showed that DAPI-bright cells were increased in size compared to WT cells (Fig. 4B). The area and minimum diameter were significantly larger in dHET sperm compared to WT sperm (Fig. 4C-D). Additionally, the hook seems more bent, leading to an increase in measured hook length in dHET sperm (Fig. 4E). Next, we evaluated nuclei sizes of testicular late-stage spermatids isolated from testis. Spermatid nuclei of dHET mice were larger compared to WT spermatids (Fig. S5A-B), showing increased nuclei area and minimum diameters compared from WT spermatids (Fig. S5C-D). Further, the ellipticity (height/width ratio) of dHET spermatids was increased compared to WT spermatids (Fig. S5E). Thus, in testes of dHET males, nuclei are less condensed suggesting impaired protamination. Nevertheless, it is important to note that late-stage spermatids did not show apparent differences in electron density in testicular transmission electron micrographs (TEM) (Fig. S5F).

**Fig. 4.**
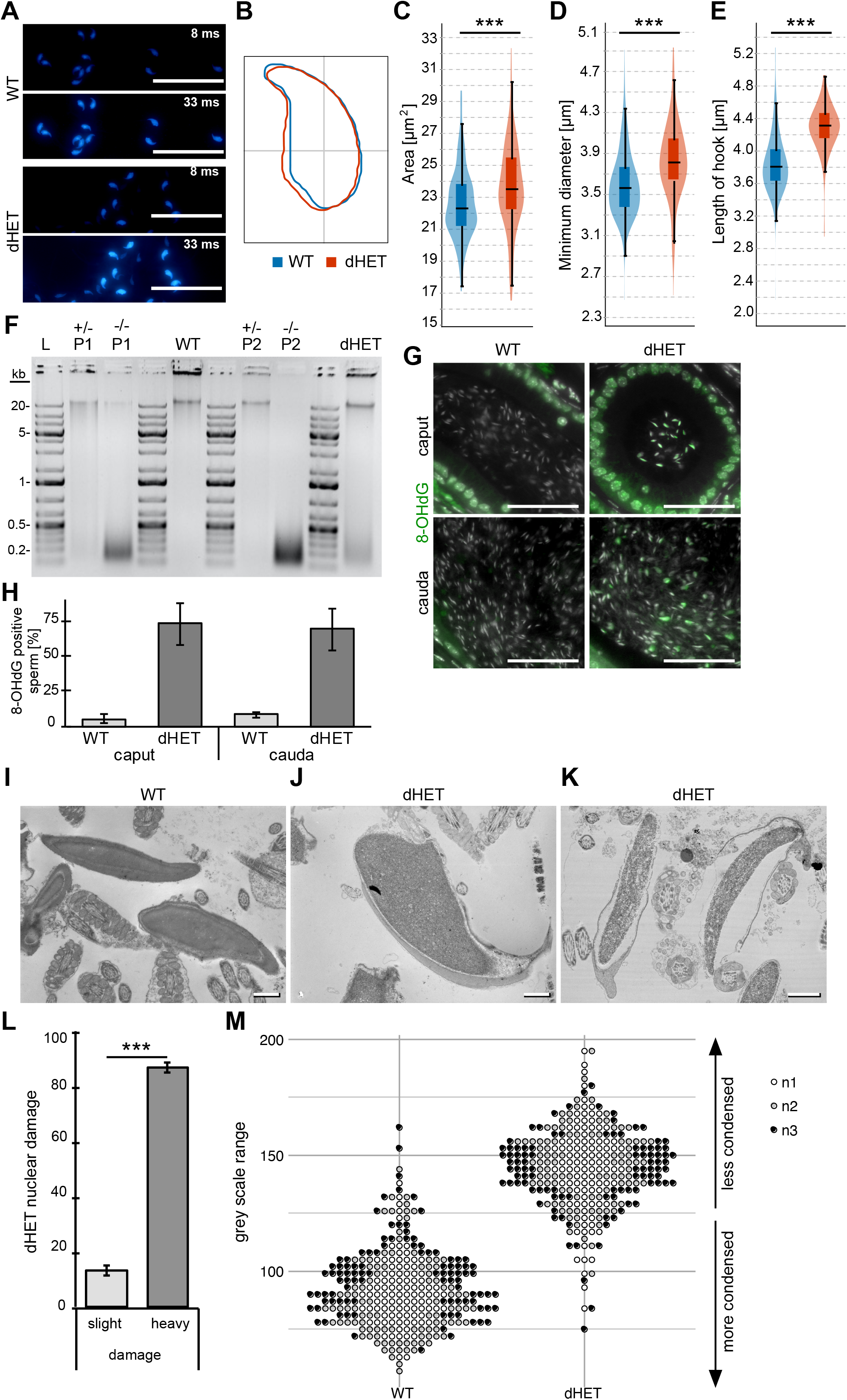
Analysis of DNA hypercondensation and damage in dHET sperm. (A) Representative images of DAPI-stained WT and dHET mature sperm. Sperm are visualized twice with different exposure times (8 ms and 33 ms), showing examples of DAPI-bright and DAPI-weak sperm in a dHET sample. Scale bars: 50 μm. (B) Sperm nuclei consensus shapes of WT and dHET epididymal sperm. Only DAPI-bright dHET cells were included (n = 3 males per genotype). (C-E) Violin plots depicting the area (C), minimum diameter (D) and length of hook (E) of WT and dHET epididymal sperm. (F) Agarose gel loaded with genomic DNA isolated from WT, *Prm1*^*+/-*^ (P1^+/-^), *Prm1*^*-/-*^ (P1^-/-^), *Prm2*^*+/-*^ (P2^+/-^), *Prm2*^*-/-*^ (P2^-/-^), and dHET cauda epididymal sperm. Part of the data has been published before ^26^. (G) IHC against 8-OHdG on epididymal (caput and cauda) tissue from WT and dHET males (Dapi counterstain in grey) Scale bars: 50 μm. (H) Percentage of 8-OHdG-positive sperm from WT and dHET sperm from caput and cauda tissue (n = 3 males per genotype). (I-K) Representative TEM images from (I) WT and (J-K) dHET mature sperm. Scale bars: 1 μm. (J) Ratio of slightly damaged (as shown in J) and heavily damaged (as shown in K) dHET sperm. Data are mean ± s.d. and were analyzed by two-tailed, unpaired Student’s t-test (***p<0.001). (M) Quantification of the difference in grey scale (grey scale range) measured for epididymal sperm nuclei from WT and dHET males (n = number of males).

Next, we isolated DNA from mature sperm to detect DNA fragmentation in *Prm1*^*+/-*^, *Prm1*^*-/-*^, *Prm2*^*+/-*^, *Prm2*^*-/-*^, dHET and WT sperm (Fig. 4F). DNA from dHET sperm was partially intact and partially fragmented. Of note, DNA from *Prm1*^*+/-*^ and *Prm2*^*+/-*^ sperm appeared mostly intact, while DNA from *Prm1*^*-/-*^ and *Prm2*^*-/-*^ samples was completely fragmented.

Since sperm of *Prm1*^*-/-*^ and *Prm2*^*-/-*^ mice have been shown to undergo ROS mediated degradation during epididymal transit ^26, 28^, we stained the ROS-mediated DNA damage marker 8-OHdG in epididymal tissue sections. In case of dHET sections, 73% and 69% cells in caput and cauda displayed 8-OHdG staining indicative for DNA damage, compared to 5% and 8% in caput and cauda of WT mice (Fig. 4G-H).

Next, we examined mature dHET sperm using TEM (Fig. 4I-M). In contrast to the results from late-stage testicular spermatids, clear differences were visible in epidydimal sperm. As seen in EN staining the vast majority of sperm presented with disrupted membranes no longer attached to the nucleus (Fig. 4K). Further, most dHET sperm presented with granulated and less contrasted nuclei, indicative of severe DNA damage compared to the electron dense WT sperm nuclei. Part of the mature sperm extracted from dHET cauda epididymis presented with seemingly intact DNA (4J, L), however the DNA appeared only partially hypercondensed and more granulated in comparison to WT sperm (Fig. 4I-J). Analysis of the greyscale within nuclei revealed that epididymal sperm from dHET mice were generally less condensed compared to WT sperm, indicative for impaired DNA hypercondensation (Fig. 4M).

### Chromatin remodeling and eviction of nuclear proteins disturbed in dHET mice

Since we had shown previously, that loss of *Prm1* or *Prm2* leads to retention of histones, we next analyzed protamination and nuclear proteins in dHET sperm in more detail.

Sperm isolated from cauda epididymis were stained with Chromomycin A3 (CMA3), an intercalating dye which does not stain protaminated DNA ^37^. In dHET mice, 99% of sperm were CMA-positive compared to 2% of sperm from WT males, which indicates insufficient protamination and supports the results from the TEM analysis (Fig. 5A-B). Of note, DAPI-weak nuclei also appeared weaker in CMA3 staining, again indicating DNA fragmentation.

**Fig. 5.**
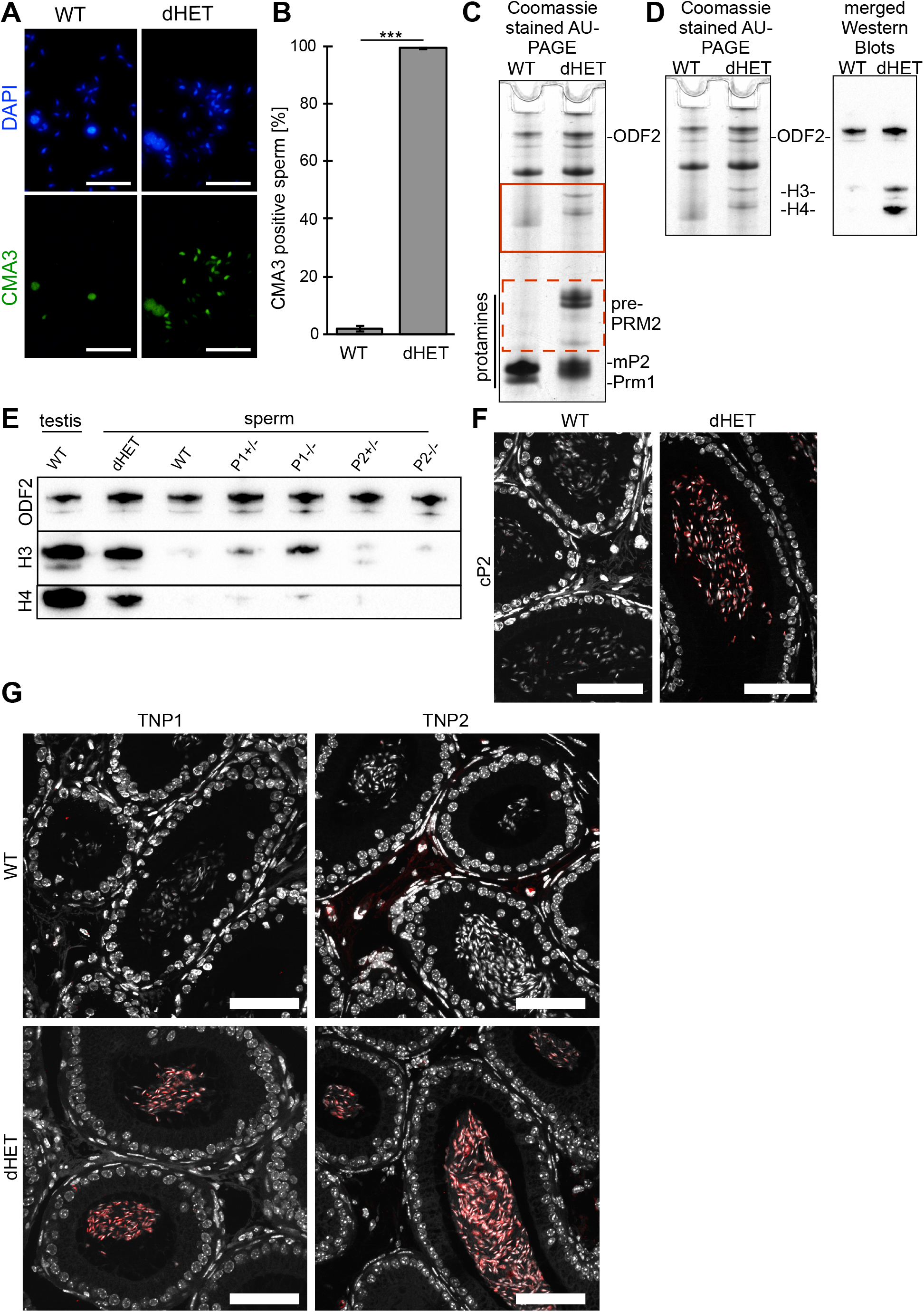
Protamination and retention of pre-PRM2, TNPs and histones in dHET sperm. (A) CMA3 staining of mature WT and dHET sperm counterstained by DAPI. Scale bars: 30 μm. (B) Quantification of CMA3-stained nuclei in WT and dHET samples. Data are mean ± s.d. and were analyzed by two-tailed, unpaired Student’s t-test (***p<0.001). (C) Representative image of a Coomassie stained AU-PAGE of basic nuclear-enriched proteins isolated from WT and dHET epididymal sperm. Bands corresponding to ODF2 and protamines are marked. Bands containing predominantly PRM1 and mature PRM2 (mP2) are labelled and areas of bands containing Prm2 precursors are marked (red dashed box). Non-protamine, nuclear-enriched proteins are found to be differentially abundant in dHET compared to WT sperm (red box). (D) Upper part of image shown in (C) compared to merged Western Blots against ODF2, histone H3 and histone H4 (pan-H3 and pan-H4 antibodies). (E) Western blots against histone H3 and H4 basic protein extractions of WT testis lysate and sperm samples form WT, dHET, *Prm1*^*+/-*^ (P1^+/-^), *Prm1*^*-/-*^ (P1^-/-^), *Prm2*^*+/-*^ (P2^+/-^), *Prm2*^*-/-*^ (P2^-/-^). ODF2 was used as loading control, as described before ^35^. (F) IHC against pre-PRM2 (antibody epitope in cP2) on WT and dHET epididymal caput tissue sections. (G) IHC against transition proteins TNP1 and TNP2 on WT and dHET epididymal caput tissue sections. Scales (F-G): 50 µm

Coomassie-stained AU-PAGE gels revealed that in dHET sperm, large amounts of pre-PRM2 were detectable (Fig. 5C, red dashed box). Further, non-protamine nuclear proteins appeared to be differentially abundant (Fig. 5C, red box). The two most prominent bands contain histones H3 and H4 (Fig. 5D).

Histone retention was further evaluated by western blot of WT, dHET, *Prm1*^*+/-*^, *Prm2*^*+/-*^, *Prm1*^*-/-*^ and *Prm2*^*-/-*^ sperm nuclear protein extracts (Fig. 5E, Fig. S6). Each lane was loaded with basic nuclear enriched protein extractions equivalent to 3 million sperm. ODF2 was used as a loading control ^26^. dHET sperm samples exhibited the highest level of histone retention followed by *Prm1*^*+/-*^ and *Prm1*^*-/-*^ sperm. In *Prm2*^*+/-*^ and *Prm2*^*-/-*^ sperm, differences in histone H3 and H4 content were not obvious compared to WT samples.

Unprocessed and/or incompletely processed PRM2 was further detected using IHC staining in caput epididymis sperm of dHET mice (Fig. 5F). Further, large amounts of TNP1 and TNP2 were retained in dHET epidydimal sperm compared to WT sperm (Fig. 5G). Altogether this suggests that in dHET males the histone-to-protamine transition is disturbed to an even larger degree as seen in *Prm1*^*-/-*^ and *Prm2*^*-/-*^ mice ^26^.

### dHET sperm are capable of fertilization, but embryonic development arrests after zygotic genome activation

Since a fraction of dHET sperm presented with intact DNA we tested fertilization capacity. Oocytes were isolated from females mated with dHET and WT males. As seen in Fig. 6A, despite the general damage detected in dHET sperm, 23% of oocytes developed to 2-cell stage indicative of fertilization (Fig. 6A). The fertilization rate, however, was significantly reduced with 23% compared to 78% in the control. In total, 106 oocytes were extracted from six females after mating with WT males and 136 oocytes from eight females mated with dHET males. In WT matings, 82 oocytes progressed to 2-cell stage embryos, of which a total of 26 developed into blastocysts. In dHET matings, 29 embryos reached the 2-cell stage. Of those, 17 embryos further developed into 4-8-cell stage (Fig. 6B). Embryonic development arrested latest at 8-cell stage.

**Fig. 6.**
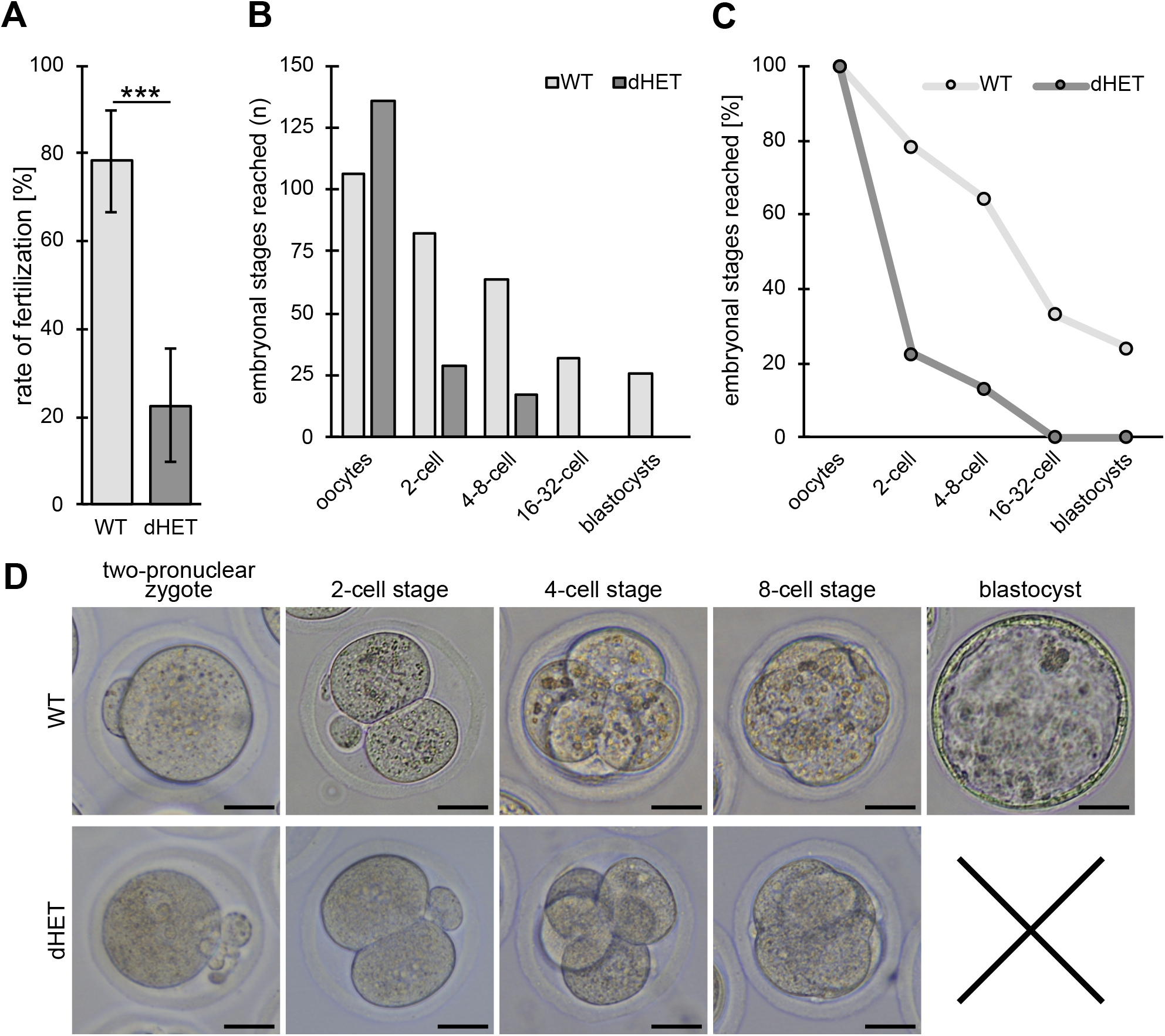
Embryonic development after breeding with dHET males. (A) Percentage of oocytes that developed into 2-cell stage in vitro. Oocytes were isolated 0.5 dpc from superovulated WT females mated with dHET and WT males. (B) Total numbers of oocytes and early embryo stages monitored. (C) Percentages of total oocytes reaching 2-cell, 4-8-cell, 16-32-cell and blastocyst stage. (D) Representative images of early-stage embryos after fertilization by WT and dHET males. (n = 6 females mated with WT and 8 females mated with dHET males). Data are mean ± s.d. and were analyzed by two-tailed, unpaired Student’s t-test (***p<0.001). Scale bars: 20 μm.

In case of dHET, 13% of the fertilized oocytes progressed to 8 cell stage, compared to 64% in case of the WT males (Fig. 6C). The embryos did not reveal any overt abnormalities in dHET samples compared to the control (Fig. 6D).

Our results show that dHET sperm are capable of fertilizing an oocyte, but embryos undergo developmental arrest at an early cleavage stage.

## Discussion

In this study, *Prm1*^+/-^*Prm2*^*+/-*^ mice were generated and analyzed. Male dHET mice were infertile with sperm showing incomplete DNA protamination, despite maintaining total PRM content and PRM ratio. Histones, TNPs and pre-PRM2 were retained in dHET sperm. Increased levels of oxidative stress induced DNA damage was apparent in epididymal sperm. Mature sperm exhibited partially fragmented DNA, membrane degradation and severely reduced motility. A fraction of dHET sperm was capable of fertilization, however resulting embryos arrested during early cleavage stages.

With our protamine mouse models, we have shown that loss of either protamine (*Prm1*^*-/-*^ and *Prm2*^*-/-*^) and loss of one allele of *Prm1* and *Prm2* each (dHET) leads to infertility in mice. *Prm2*^*+/-*^ males were fertile, while *Prm1*^*+/-*^ males were subfertile, suggesting that loss of one *Prm* allele can be tolerated ^26-28^. Hence, loss of two *Prm* alleles is detrimental for male fertility.

We found, that in epidydimal sperm from dHET males the protamine ratio was not altered. In previous studies, aberrations of the protamine ratio have been correlated to sub- and infertility both in humans and mice ^15, 18, 19, 24, 38, 39^. Epidydimal sperm from *Prm1*^*+/-*^ mice showed a shift of the protamine ratio to 1:5 and were subfertile. Noteworthy, *Prm2*^*+/-*^ males, which show a protamine ratio comparable to WT, were fertile ^26^. So, *Prm1*^*+/-*^ males display shifted ratio, while the ratio in dHET males remains unaffected and comparable to WT. Thus, it appears, that loss of one *Prm1* allele generates a transcriptional imbalance which results in overrepresentation of PRM2 which is not the case if one allele of both *Prm1* and *Prm*2 is deleted. Additionally, in dHET mice protein levels of PRMs are not reduced. This demonstrates that there is no gene-dosage effect in case of dHET.

In dHET epididymal sperm we found increased levels of pre-PRM2 and decreased levels of mP2. Increased levels of pre-PRM2 were also observed in *Prm1*^*+/-*^ and *Prm2*^*+/-*^ male mice ^26^. Approximately 55% of total PRM2 was pre-PRM2 in *Prm1*^*+/-*^ sperm and 35% in *Prm2*^*+/-*^ sperm^26^. In *Prm1*^*+/-*^ and *Prm2*^*+/-*^ sperm, the relative levels of mP2 to total PRM were, however, not significantly different from WT. In contrast, the relative levels of mP2 to total PRM were significantly reduced in dHET sperm. Hence, in dHET sperm PRM2 processing appears to be impaired to a larger extent compared to *Prm1*^*+/-*^ and *Prm2*^*+/-*^ sperm.

Of note, retained PRM2 precursors were also detected in mice lacking one or both TNPs or the sperm-specific histone variant H2A.L.2 ^2, 40, 41^. In infertile patients, increased levels of PRM2 precursors have been negatively correlated to sperm count, normal sperm morphology and motility ^24^. Further, increased levels of PRM2 precursors were correlated with increased percentage of TUNEL positive spermatozoa. Moreover, Rezaei-Gazik *et al*. found that elevated PRM2 precursor levels in sperm correlated with decreased ICSI success ^42^. This implies, that the presence of PRM2 precursors in mature sperm interferes with fertility. We speculate that at least some of the patients might present with variants (mutations) affecting PRM1 and PRM2 genes.

In view of these results, the question arises whether deletion of the cP2 domain and hence skipping of PRM2 processing could ameliorate incomplete histone-to-protamine transition. However, a mouse genetically modified to express only mature PRM2 without cP2 (*Prm2*^*+/1c*^, *Prm2*^*-/Δc*^) rendered males infertile indicating the requirement of processing of pre-PRM2 ^35^. Of note, *Prm2*^*+/1c*^ sperm show low levels of retained pre-PRM2 as well. Hence, PRM2 processing is essential ensure proper histone-to-protamine transition.

In dHET male mice CMA3 staining revealed, that 99% of epididymal sperm displays insufficient protamination. In idiopathic oligoasthenoteratozoospermic men increased CMA3 staining of sperm is negatively correlated to sperm concentration, motility and normal sperm morphology ^43^. Notably, 98% of *Prm1*^*+/-*^ sperm, but only 29% of *Prm2*^*+/-*^ epididymal sperm were CMA3 positive. This argues that loss of one allele of *Prm1* alone seems to cause the high degree of non protaminated DNA.

Further, we observed retention of histones H3 and H4 in dHET sperm to a larger degree compared to *Prm1*^*-/-*^ or *Prm2*^*-/-*^ sperm. Large amounts of retained histones and TNPs correlate to increased CMA3 staining. This could help to explain why epididymal sperm of dHET mice appeared less condensed in TEM. Increased levels of histone and TNP retention have been shown to be a result of incomplete histone-to-protamine transition ^26-28, 35, 44-47^, contributing to infertility in dHET.

Additional to PRM2 precursor bands visible in WT testis samples, prominent bands in the protamine region of AU gels were detected for dHET sperm and testis samples. These were not or only weakly visible in WT samples and might represent immature PRM2 proteoforms and other PRM2 or PRM1 proteoforms. A recent study defined protamine proteoforms in human sperm using a refined top-down mass spectrometry approach ^48^. Sperm presenting with shifted PRM ratio showed an accumulation of specific immature PRM2 proteoforms compared to sperm from normozoospermic men. Additionally, sperm from obese men and men of advanced age showed a loss of specific P1 proteoforms. Identification and analysis of these PRM proteoforms forms in dHET sperm could help to explain the consequences on PRM2 processing and posttranslational modifications of PRMs in more detail.

dHET sperm showed various damages, including fragmented DNA, disrupted membranes, acrosome and tail defects. Our group has shown before that protamine-deficient and -modified mice show ROS-mediated sperm damage in epididymides ^26, 28, 35^. Apparently, a ROS-mediated destruction cascade is initiated during epididymal transit, causing the gradual accumulation of sperm damage ^28^. Indeed, it has been shown that once ROS levels exceed scavenging levels, lipid aldehydes target mitochondrial proteins, leading to yet more ROS production and a self-perpetuating cycle, which finally causes severe sperm damage and fertility problems ^49^. While 69% cauda epididymal sperm from dHET sperm were positive for 8-OhdG, 64% of *Prm1*^*-/-*^ sperm were 8-OhdG positive. In comparison, only 3% of *Prm1*^*+/-*^ sperm stained 8-OhdG positive, while no significant differences were detected in *Prm2*^*+/-*^ sperm compared WT ^26, 28^. DNA fragmentation visualized on agarose gel was higher in dHET sperm compared to *Prm1*^*+/-*^ and *Prm2*^*+/-*^ sperm, but lower compared to *Prm1*^*-/-*^ and *Prm2*^*-/-*^ sperm, which appeared completely fragmented. This suggests, that loss of two protamine alleles increases the sperm susceptibility to ROS-mediated damage.

Interestingly, in dHET mature sperm presents as a DAPI-bright and DAPI-weak. DAPI-bright dHET sperm resembled WT, *Prm1*^*+/-*^ and *Prm2*^*+/-*^ sperm. The DAPI-weak sperm nuclei strongly resemble *Prm1*^*-/-*^ sperm nuclei, which presented with fragmented DNA ^26^. Hence, we hypothesize, that a fraction of the dHET sperm undergoes DNA degradation during epididymal transit resulting in DAPI-weak smaller nuclei. The fact, that on an agarose gel DNA from dHET sperm presented either as high molecular weight, intact or degraded at 200-500bp further supports this hypothesis. Likewise, in TEM some sperm nuclei from cauda epididymis of dHET males appeared less condensed with clear vacuoles indicative of chromatin damage.

How can these apparent two populations of sperm be explained? Haploid dHET sperm are either *Prm1-/Prm2-* or *Prm1+/Prm2+* giving a rationale for the effect. However, our IHC showed that all dHET sperm contain PRM1 and PRM2 protein. This is due to transcript sharing between developing spermatids. In mice, heterozygous for two Robertsonian translocations involving chromosome 16 including the *Prm1* locus, it has been shown that *Prm1* transcripts, are shared within a syncytium resulting in an equal distribution of transcripts ^50^.

Either the pathophysiological differences within dHET sperm originate from a so far not detected mechanism, or the methods used to detect and quantify amounts of protamine protein might not be sensitive enough to detect subtle differences leading to the observed defects. Only 9% of dHET sperm were motile compared to 71% of WT sperm. In comparison, approx. 23% of *Prm1*^*+/-*^ sperm were motile, while *Prm2*^*+/-*^ sperm showed no significant reduction in motility compared to WT sperm. Surprisingly, in contrast to sperm from *Prm1*^*-/-*^ and *Prm2*^*-/-*^ males, a small fraction of dHET sperm is capable of fertilization. Compared to the WT control (78%), fertilization rate of dHET males was significantly reduced (23%). Using dHET sperm, embryonic development did not proceed past the 8-cell stage. We demonstrate, that sperm from dHET males display large amounts of retained pre-PRM2, histones H3 and H4, as well as both TNPs. We hypothesize, that this interferes with PRM to histone transition and zygotic genome activation. Incorrect histone modification patterns might also heavily disturb the zygotic transcriptome, since this epigenetic information is known to strongly affect gene expression ^51^. DNA damage could be another reason for the developmental arrest. It has been shown that DNA damage in the paternal chromatin can lead to embryonic arrest at later points than the 2-cell stage ^52^. As shown by CMA3 staining and nuclei grey scale measurement in TEM, the whole dHET sperm population (including the DAPI-bright sperm) displays DNA compaction defects. Overall, dHET sperm are capable of fertilization, allowing to study the effects of defective protamination on early embryonic development. This will be subject to further studies.

In summary, we showed that dHET male mice are infertile and sperm present with retention of pre-PRM2, histones and TNPs leading to less condensed DNA triggering a ROS-mediated destruction cascade during epididymal transit. This led to DNA damage and a wide variety of secondary defects in mature sperm including membrane disruption, decreased motility and degradation of the acrosome and disturbance of the axonemal structures.

Our study describes a mouse model with protamine-mediated fertility problems that cannot be explained by alterations of the PRM ratio. Our results indicate that measurement of the PRM1/PRM2 ratio is not sufficient to predict fertility outcomes. In consequence, we hypothesize that male factor infertility patients would benefit from CMA3 diagnosis with additional PRM2 precursor screening.

## Acknowledgements

We thank Gaby Beine, Greta Zech, Andrea Jäger and Angela Egert for excellent technical assistance. We thank the Microscopy Core Facility of the Medical Faculty at the University of Bonn for providing support and instrumentation funded by the DFG (388171357).

## Author contributions

Conceptualization: G.E.M., H.S., L.A.; Methodology: G.E.M., S.S., L.A.; Formal analysis: G.E.M., C.W., L.A.; Investigation: G.E.M., C.W., M.D.L., A.K., L.A.; Writing - original draft:

G.E.M., C.W.; Writing - review & editing: G.E.M., C.W., L.A., H.S.; Visualization: G.E.M., C.W.; Supervision: H.S., L.A.; Project administration: H.S.; Funding acquisition: H.S.

