## Supplementary_Data for "Male mice heterozygous for Protamine-1 and Protamine-2 are infertile displaying sperm damage and retention of Protamine-2 precursors, transition proteins and histones"

### Figure S1

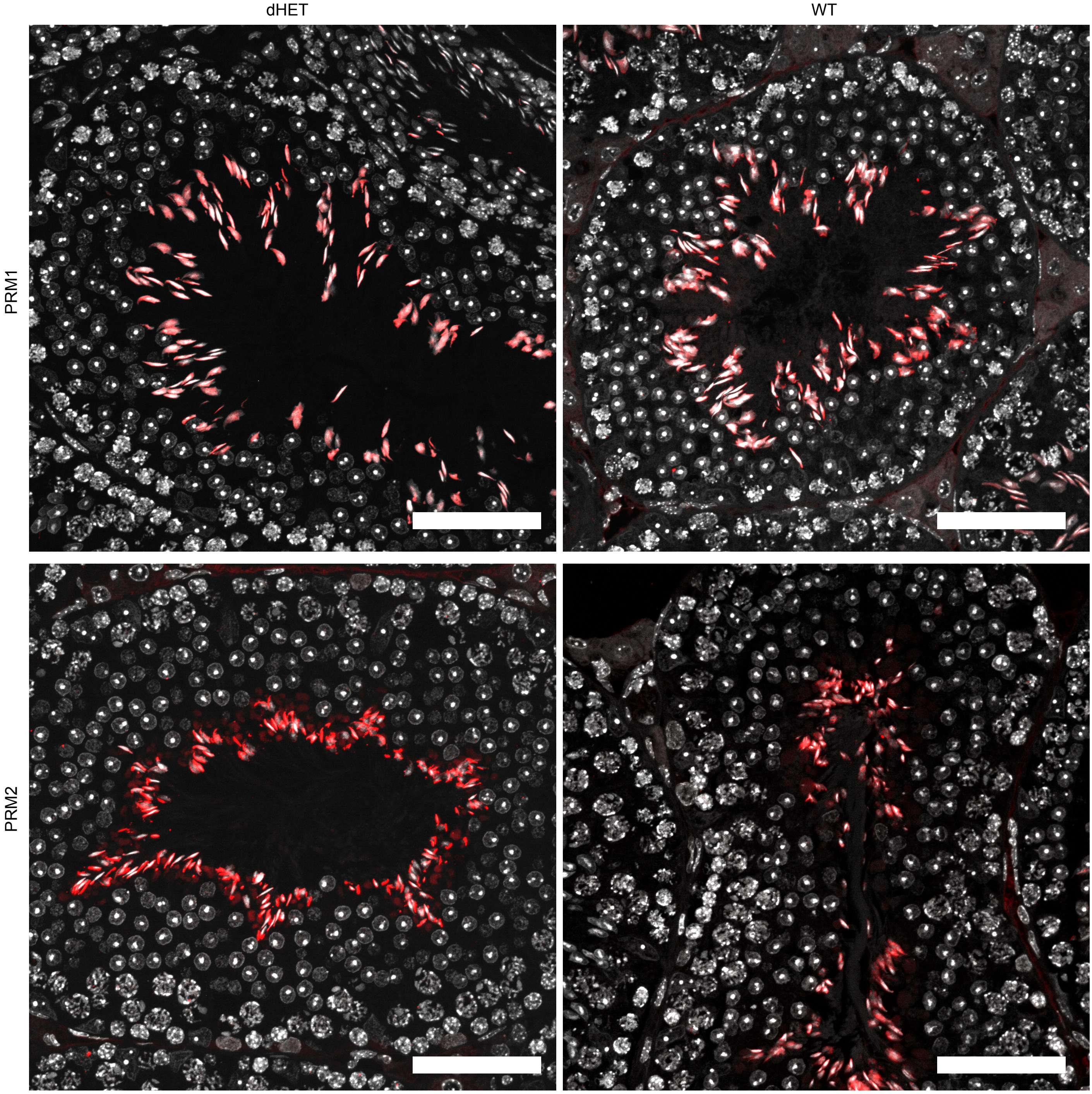


**Fig. S1. Fluorescent Immunohistochemical (IHS) staining against PRM1 and PRM2.** IHC staining of PRM1 and PRM2 (mP2-targeting antibody) on dHET and WT testis sections. Scale bars: 50 µm.

### Figure S2

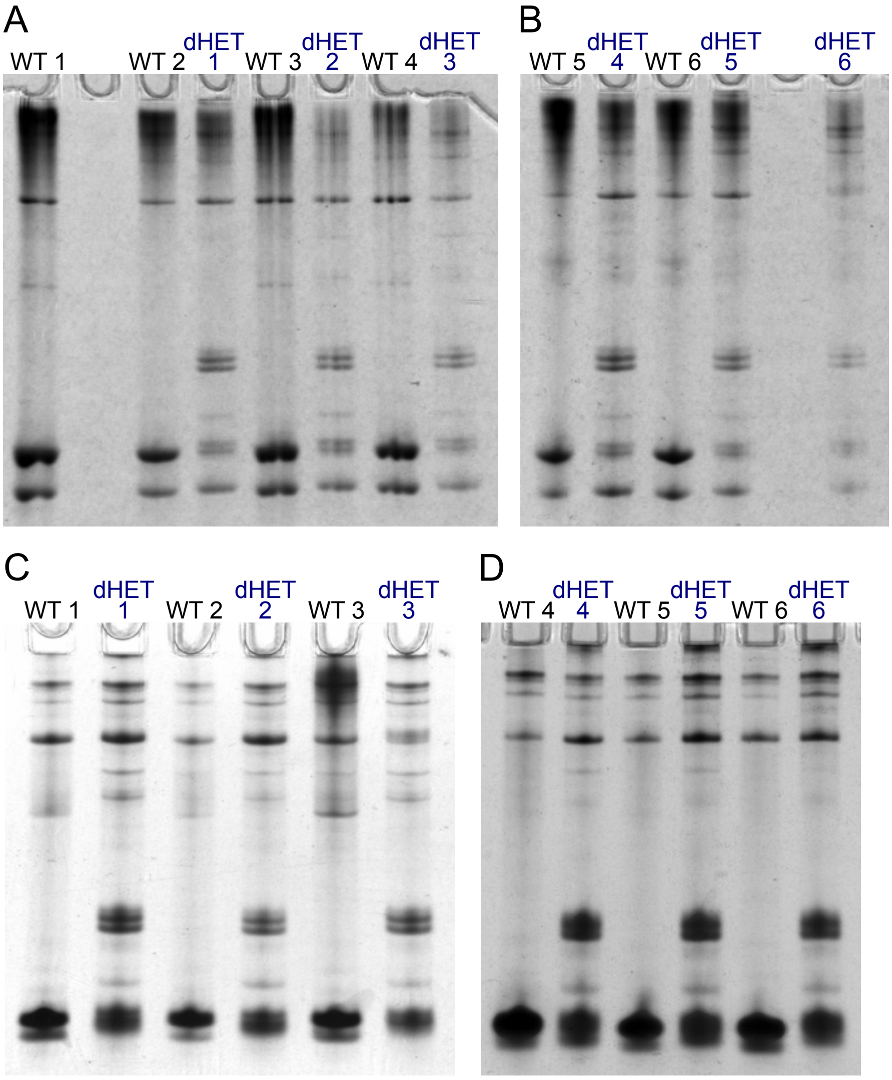


**Fig. S2. Thiourea and AU gels used for analysis of basic protein extractions from epididymal sperm.** (A, B) Thiourea gels depicting protein extractions from six males per genotype. Primarily used for separation of protamine bands. (C, D) Acetic acid-urea gels showing protein extractions from six males per genotype. Primarily used for separation and visualization of histone bands.

#### Figure S3

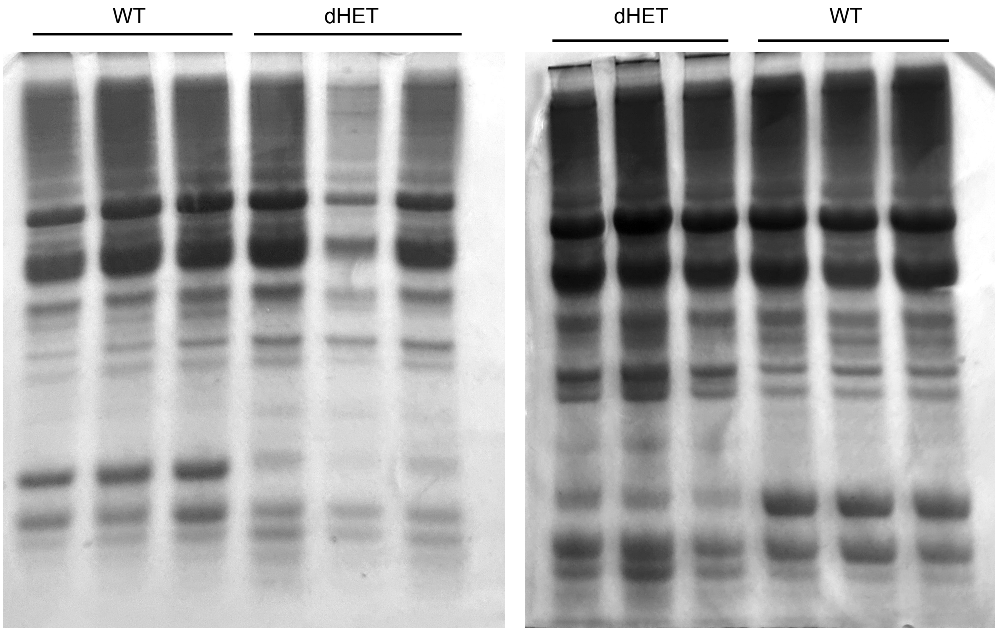


**Fig. S3. Thiourea gels used for analysis of basic protein extractions from testis.** Thiourea gels depicting protein extractions from six males per genotype.

### Figure S4

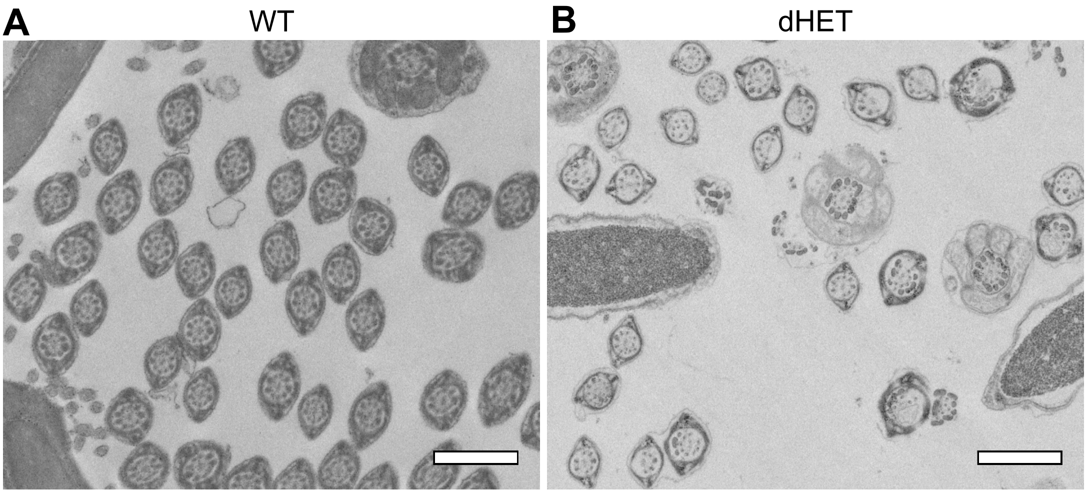


**Fig. S4. Transmission electron micrographs of flagella cross sections of dHET and WT sperm.** (A) TEM flagella cross sections of WT sperm isolated from cauda epididymis. (B) TEM flagella cross sections of dHET sperm isolated from cauda epididymis showing secondary damage of 9+2 axonemal structure. Scale bars: 1 µm.

#### Figure S5

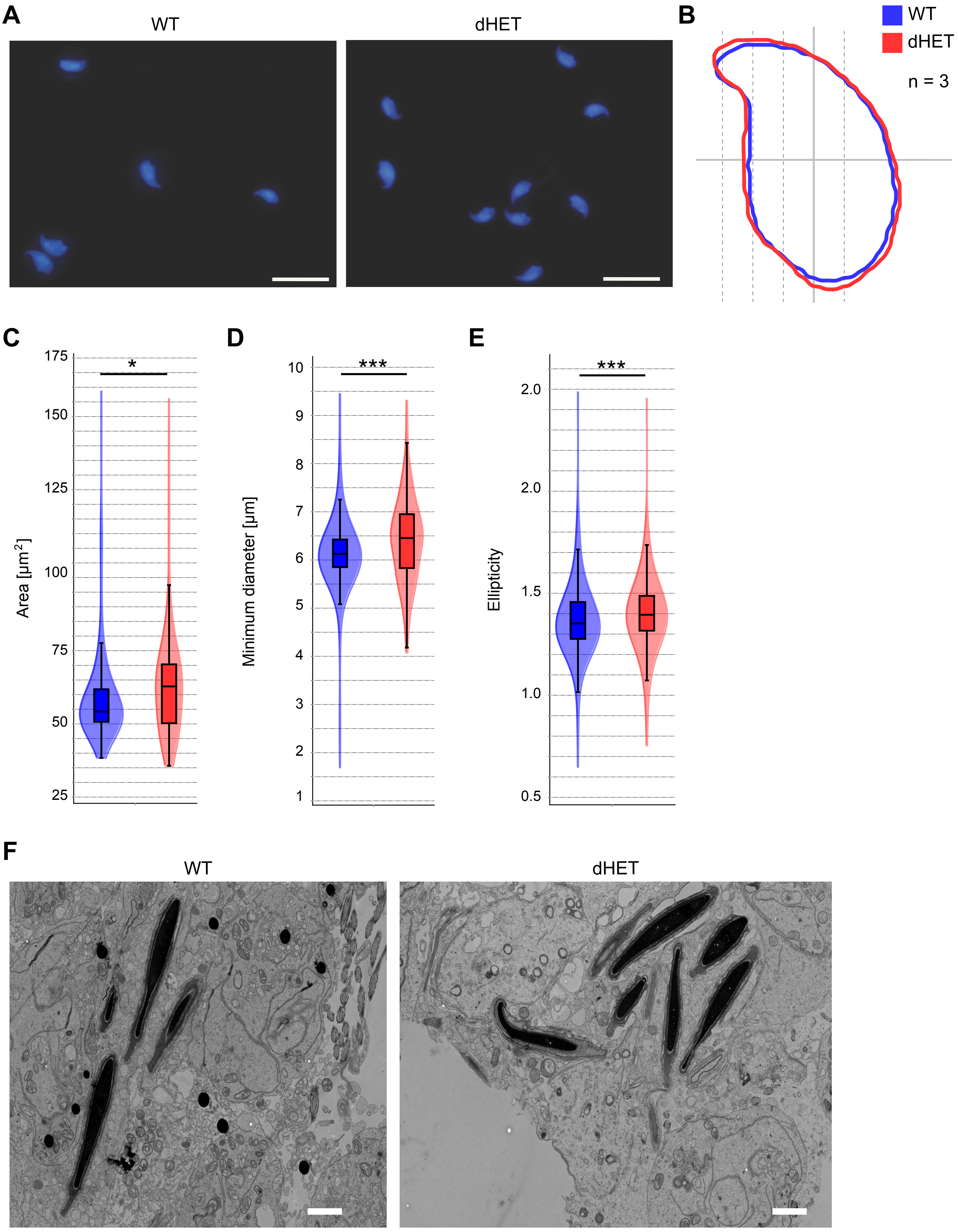


**Fig. S5. Shape and chromatin remodeling of dHET spermatids.** (A) Representative images of DAPI stained testicular dHET and WT spermatids (step 15-16). Scale bars: 20 µm. (B) Consensus sperm nuclei shapes of dHET and WT late-stage spermatids. (n = number of males used) A minimum of 250 spermatids per male were analyzed. (C-E) Violin plots depicting the area (C), minimum diameter (D) and ellipticity (E) of WT and dHET late-stage spermatids. (F) Transmission electron micrographs of dHET and WT testis tissue. Depicted are late-stage spermatids close to the lumen. Scale bars: 2 µm

#### Figure S6

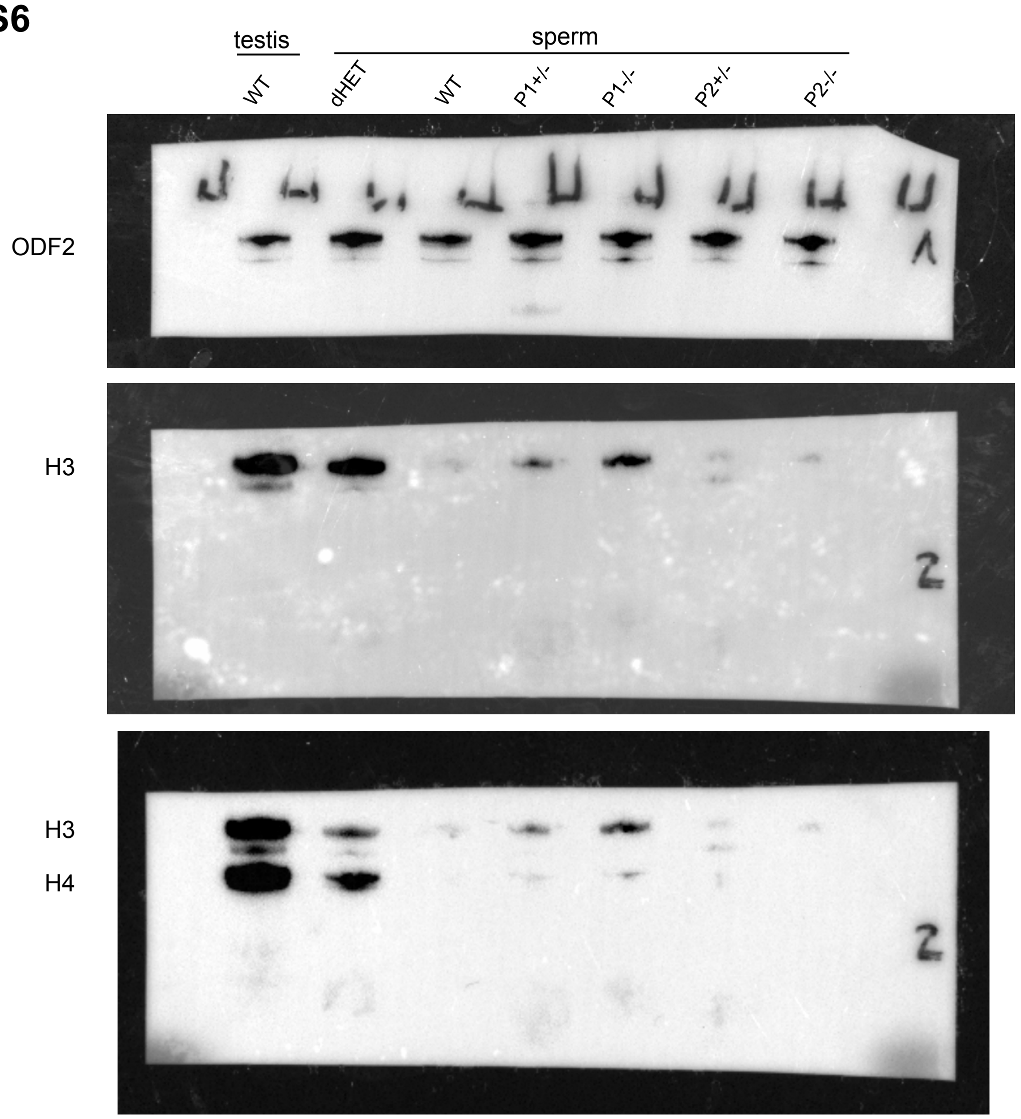


**Fig. S6. Representative Western blots against histones H3 and H4.** Protein extractions of WT testis lysate and sperm samples form WT, dHET, *Prm1+/-* (P1+/-), *Prm1-/-* (P1-/-), *Prm2+/-* (P2+/-), *Prm2-/-* (P2-/-) mice were used. The membrane was processed in two parts. The upper half (1) was used to detect ODF2. The lower half (2) was used to detect H3, followed by H4.

**Table S1**

| Fertility analysis *Prm1△139+/-* male mice | | | | | | | | |
|---|---|---|---|---|---|---|---|---|
| mouse # | genotype | litter sizes | | | litters | average litter size | pregnancy rate | plugs monitored |
| 7953 | +/- | | | | 0 | 0 | 0 | 6 |
| 7952 | +/- | 1 | | | 1 | 1 | 0.2 | 5 |
| 5705 | +/- | 2 | 4 | 6 | 3 | 4 | 0.6 | 5 |
| 5703 | +/- | 1 | | | 1 | 1 | 0.2 | 5 |
| 7951 | +/- | | | | 0 | 0 | 0 | 5 |
| 5994 | +/- | | | | 0 | 0 | 0 | 5 |
